# Additional blood meals increase mosquito infectivity by accelerating the transcriptional development of *Plasmodium* sporozoites

**DOI:** 10.64898/2026.09.26.754681

**Authors:** W. Robert Shaw, Philipp Schwabl, Maurice A. Itoe, Shriya Anandjee, Jamie Kauffman, Lisa H. Verzier, Yan Yan, Daniel E. Neafsey, Flaminia Catteruccia

## Abstract

Sporozoites, the infectious stage of *Plasmodium* malaria parasites, must invade two different tissues in two very different hosts to ensure transmission: the mosquito salivary glands and the human liver. Whether infectivity to hepatocytes is affected by physiological components, such as the availability of nutrients within the *Anopheles* female, is not well understood. Here we show that *Plasmodium falciparum* sporozoite infectivity to hepatocytes is increased if mosquitoes take an additional blood meal while oocysts are developing. Infection rates of HC04.J7 hepatocytes at 24 h were 1.8-fold higher for *P. falciparum* salivary gland sporozoites isolated from *Anopheles gambiae* females fed twice (2BF) compared to those which only received the infectious blood meal (1BF). This increase in infectivity was observed at an early infection timepoint (day 11) and was accompanied by a marked shift in the transcriptional profiles of 2BF salivary gland sporozoites, identifying genes likely involved in infection and development in the mammalian host. Overall, our data reveal new factors potentially involved in sporozoite infectivity, and have broad implications for understanding transmission of malaria in endemic areas where multiple blood meals commonly shape available resources for adult female mosquitoes.

## Introduction

Malaria continues to be a devastating disease in tropical and sub-tropical regions, with a global annual burden of 282 million cases and 610,000 deaths reported for 2024 (1). The disease is caused by apicomplexan parasites of the *Plasmodium* genus, with *Plasmodium falciparum* accounting for >95% of deaths (1). The parasite transmission cycle begins when an *Anopheles* female ingests gametocytes during a bloodmeal to develop eggs. During a lengthy developmental process, parasites transform into oocysts that encyst under the basal lamina of the midgut epithelium, where they undergo multiple rounds of DNA replication without division. After 7–10 days, the oocyst segments into thousands of daughter sporozoites, which then egress from the oocyst in the following days and invade the mosquito salivary glands for transmission to the next human host. The time taken for parasites to reach an infective state within the salivary glands after initial mosquito uptake is known as the extrinsic incubation period (EIP) and is a key parameter in models of malaria transmission, due to the short (2-3 weeks) lifespan of female mosquitoes (2). During this time, *Anopheles* females take several blood meals in order to undergo several rounds of egg development and egg laying (2). With others, we have shown that the EIP is accelerated by an additional blood meal, causing the earlier appearance of sporozoites in the salivary glands. Multiple blood feeding behavior, which occurs frequently in field settings, is therefore epidemiologically important as it causes younger mosquitoes to also contribute towards transmission in sub-Saharan Africa (3). However, a full understanding of how mosquito feeding behavior and physiology surrounding blood meal acquisition and reproduction impact malaria transmission is still lacking, although this knowledge is key to optimizing the effective deployment of control interventions such as long-lasting insecticide-treated nets and indoor residual insecticidal sprays.

One of the outstanding questions is whether, aside from accelerating the occurrence of salivary gland invasion, an additional blood meal affects sporozoite infectivity to liver cells. In the course of an infection, sporozoites released from mature oocysts increase their infectivity as they travel through the open circulation of the mosquito (hemolymph) and invade the salivary glands (4–6). Once across the basal lamina, sporozoites invade the secretory acinar cells and access the lumen of the gland duct, at which point they are thought to enter a stage of translational quiescence until transmission to the next host (7–13). Previous work in the rodent and avian model malaria species *Plasmodium berghei* and *Plasmodium gallinaceum*, respectively, has shown that the sporozoite infectivity rate changes over time in the salivary glands, gradually increasing between 12–18 days (d) post infectious blood meal (pIBM) before decreasing over several weeks (4, 5, 14, 15). This acquisition of infectivity over time suggests that either translational quiescence is not immediately or fully established, or that there are additional transcriptional maturation processes ongoing within the salivary glands.

The molecular nature of the acquisition of infectivity has previously been studied comparing RNA-seq and single-cell RNA-seq (scRNA-seq) datasets of sporozoites derived from midgut oocysts, the hemolymph, and the salivary glands (16–23). These studies revealed a shift in sporozoite transcription that is coincident with their developmental journey through the mosquito, with midgut sporozoites expressing higher levels of transcripts required for salivary gland invasion, while salivary gland sporozoites express transcripts required for subsequent liver-stage development. Comparisons to proteomic datasets and genetic mutants also revealed extensive translational control of transcript expression (10–13, 18, 19). Many of these publications focused on late timepoints for salivary gland sporozoites collection (≥16 d pIBM) yet found a large amount of transcriptional variation within populations (in both *P. falciparum* and *P. vivax*) (22, 23). This suggests that, even after an extended period of time within the salivary glands, a proportion of sporozoites may not be entirely quiescent or mature. The consequences of this transcriptional heterogeneity, however, remain unclear.

Here we assess the impact of an additional blood meal on sporozoite heterogeneity, maturity and infectivity using scRNA-seq and *in vitro* hepatocyte infection assays. We find that at an early timepoint in salivary gland infection (11 d pIBM), sporozoites developing in females fed an additional time (2BF) are more infectious to hepatocytes than sporozoites developing in females fed only once (1BF). Through scRNA-seq at this early timepoint, we find that 2BF sporozoites are shifted forward along a maturation gradient, upregulating critical liver-stage genes. We additionally identify novel factors with potential roles in infectivity that can inform future mechanistic or vaccine studies. Our work highlights that natural mosquito blood-feeding behavior has large implications for the effective sporozoite infectious reservoir and on the likelihood of parasite transmission from mosquitoes to humans, critically informing malaria control programs.

## Methods

### Mosquito rearing

*Anopheles gambiae* (wild-type G3 strain) were reared in cages at 27 °C, 70–80 % humidity, and on a 12:12 h light:dark cycle. Adults were fed 10 % glucose *ad libitum* and weekly on purchased human blood (Research Blood Components) for colony maintenance.

### Plasmodium infections of mosquitoes

*P. falciparum* infections and additional blood feedings were carried out as in (3). Briefly, P*. falciparum* (NF54 strain) parasites were maintained as asexual stages in human erythrocytes and gametocytes were induced as previously described (24). Gametocyte-containing blood cultures were fed to cages of 4-day-old mated female mosquitoes that were then maintained in a custom-built infection glove box. After blood feeding, incompletely fed mosquitoes were removed, and water and 10 % glucose solution were provided. At 2 d pIBM, mosquitoes were provided with an oviposition site and starved to encourage blood intake at the second blood meal. At 3 d pIBM, some females were fed a second time with uninfectious blood. As above, after blood feeding, incompletely fed mosquitoes were removed, and water and 10 % glucose solution were provided to all groups. At 7 d pIBM, oocysts were imaged using a 3.2 megapixel SC30 camera on an inverted Olympus CKX41 at 10X magnification. Oocysts were quantified and sized using FIJI (25) and an automated oocyst counting program (26). As oocyst sizes within each midgut are not independent, mean oocyst sizes per midgut were calculated. At 11 and 15 d pIBM sporozoites were harvested from 12–36 mosquitoes at noon. Six replicates were performed: three replicates (1–3) compared unmanipulated 1BF mosquitoes to ds*GFP*-injected 2BF mosquitoes, which served as controls in a parallel experiment not reported here; three replicates (4–6) compared unmanipulated 1BF mosquitoes to unmanipulated 2BF mosquitoes. GFP fragments were amplified from plasmid pCR2.1-eGFP plasmid as in (3). PCR products were confirmed by gel electrophoresis and dsRNA was transcribed using the Megascript T7 transcription kit (Thermo Fisher Scientific). 690 ng of ds*GFP* dsRNA was injected (Nanoject III, Drummond) into 1-day-old females at a concentration of 10 ng/nl. As *gfp* was not expressed in these mosquitoes, gene knockdown was not determined. Effects of an additional blood meal on infection outcomes were extremely similar between these two experimental designs (**Figure S1**). While we endeavored to perform scRNA-seq and hepatocyte infection at all replicates, sporozoite intensities did not always permit parallel experiments: as such, scRNA-seq data was collected in replicates 1–3, 5 and 6; hepatocyte infection data was collected in replicates 2–4 and 6 (11 d pIBM only). Infection data were pooled between ds*GFP*-injected and unmanipulated groups and compared using Fisher’s exact (prevalence), Unpaired t (oocyst intensity), and Welch’s t (oocyst size, sporozoite intensity) tests, as appropriate (GraphPad Prism 10).

### Sporozoite isolation and purification

Mosquitoes were decapitated, surface-sterilized in 70 % ethanol for 5 s, and washed in ice-cold RPMI media (Corning) supplemented with 30 U/ml Penicillin-Streptomycin (3 %PS) (Thermo-Fisher Scientific). Salivary glands were expelled from the thorax with gentle pressure and collected into ∼100 µl RPMI+3 %PS. Sporozoites were released from salivary glands by gentle disruption using rotation of a handheld sterile pestle for 1 min. Sporozoites were purified away from mosquito debris with a 30 µm cell strainer (PluriSelect) an 17 % Accudenz gradient using a previously published protocol (27), with a modification whereby sporozoites were more gently and effectively pelleted by diluting 300 µl of the collected interphase with 700 µl RPMI+3 %PS at 4 °C, followed by centrifugation at 2000 *g*, 4 °C for 10 min. Sporozoites were quantified using a disposable hemocytometer (InCyto).

### Plasmodium infection of hepatocytes

The HC04.J7 hepatocyte cell line (28) was cultured at 37 °C, 5 % CO_2_, ambient O_2_, in complete DMEM12 media, supplemented with 10 % FBS. Thirty-five thousand cells were seeded onto collagen-coated glass coverslips in 24-well plates 24 h prior to sporozoite infection. Ten thousand purified sporozoites were added to each well in 500 µl complete DMEM12 (10 % FBS) +3 %PS and incubated for 24 h. After 24 h, cells were fixed in PFA for 10 min and outside-inside stained with 1/300 mouse monoclonal α-*Pf*CSP (clone 2A10, MRA-183A, www.beiresources.org) (29). Briefly, parasites outside cells were detected with Alexa-568 goat α-mouse, and following permeabilization with 0.1 % Triton X-100, all parasites were detected with Alexa-488 goat α-mouse. DAPI (1 ng/µl) was added to the final PBS washes. Coverslips were inverted onto glass slides in Fluoromount-G (Thermo Fisher Scientific) and imaged using a 20X air objective on a fluorescent AxioObserver microscope. Hepatocytes were counted automatically using a FIJI (25) macro to identify nuclei, and parasite invasion was confirmed manually by Alexa-488 staining alone in proximity to hepatocyte nuclei. Data were pooled between ds*GFP*-injected and unmanipulated 2BF groups, and compared to the 1BF group with an Unpaired t or Welch’s t test as appropriate.

### scRNA-seq library preparation and sequencing

We prepared single-cell libraries using the Chromium Next GEM Single Cell 3ʹ v3.1(Dual Index) system from 10x Genomics. We applied 3.3 µl cell suspension (1000 sporozoites/µl, adjusted with RPMI) per sample for the initial emulsification and reverse transcription step, followed by 12 PCR cycles to amplify resultant cell-barcoded cDNA, and another 12 PCR cycles to index samples using Dual Index TT Set A. Details on all library preparation steps are provided in 10x Genomics guide CG000315. We normalized library concentrations using qPCR (KAPA Library Quantification Kit) and submitted libraries to the Broad Institute for paired-end sequencing on the Illumina NovaSeq S4 platform.

### Read alignment, expression matrix filtration, and dimensionality reduction

We used STAR (version 2.7.2a) called by cellranger count (version 6.0.1, default parameters) to map reads to the *P. falciparum* 3D7 genome (Release 58 from https://plasmodb.org) and to generate unique molecular index (UMI) matrices summarizing gene expression per cell. An average of 33.1 million reads mapped confidently to the *P. falciparum* transcriptome (**Table S1**). We imported expression matrices into R (version 4.2.2) and performed quality control, keeping cells with: gene count ≥ 125; UMI count ≥ 400; log_10_(gene count/UMI count) > 0.80; and mitochondrial gene representation < 0.025 (**Figure S2**). We then equalized sample sizes per treatment by randomly removing cells from larger libraries within each replicate. We normalized counts using the NormalizeData, FindVariableFeatures, ScaleData workflow in Seurat (version 5.0.3) and then applied RPCA integration to the PCA reductions of each normalized dataset using Seurat’s IntegrateLayers function with k.weight set to 64. We used the first nine dimensions of the integrated PCA reduction for subsequent UMAP projection. To maximize anchoring points for integration, data from all the scRNA-seq experiments (replicates 1–3, 5 and 6) were pooled.

### Clustering and pseudotime analysis

We used the cluster_cells function in Slingshot (version 2.6.0) to assign cells to Leiden clusters (resolution = 0.00015) and to assign pseudotime values based on minimum spanning tree (getLineages) and simultaneous principal curve construction (getCurves) within the integrated UMAP space. We used Seurat’s FindConservedMarkers function to identify cluster markers genes for which fold-change averaged > 1.5 (i.e. > 50 % gene expression in cluster member vs. non-member cells) across all day 11 libraries and for which each library p-value was below 0.001. We tested for differential cell abundance between treatments using quasi-likelihood methods (glmQLFtest) in edgeR (version 3.40.2) as well as with traditional Chi-squared tests. To identify genes exhibiting differential expression with respect to UMAP trajectories, we filtered genes for spatial autocorrelation (Moran’s I > 0.20 and q-value < 10^-6^) using reversed graph embedding (learn_graph and graph_test) in Monocle3 (version 1.3.1). We plotted autocorrelated expression profiles by applying Seurat’s DoHeatmap function to Slingshot pseudotime-ordered cells exhibiting > 500 UMIs sampled randomly from 1BF and 2BF groups (1000 cells each). We also tested for pseudotime differences between treatments by Wilcoxon rank sum test. Clustering and pseudotime analyses pooled data from all scRNA-seq experiments (replicates 1-3, 5 and 6) and were highly similar to plots generated after focusing on replicates with unmanipulated mosquitoes (**Figure S3**).

### Differential expression by treatment

We identified differential gene expression between treatments by Wilcoxon rank sum analysis applied to normalized counts and blocked by replicate using the findMarkers function in scran (version 1.26.2). Here, scRNA-seq downstream analyses were focused on replicates 5 and 6 with field-like infection intensities, although gene lists were highly similar when all scRNA-seq replicates were included (**Table S2**). We applied findMarkers to the whole dataset with a fold change threshold of 1.2 and an FDR-adjusted p-value of 0.0005, as well as to cluster-defined cell populations, when, with fewer cells, the FDR-adjusted p-value was relaxed to 0.05. Leiden cluster cell assignments were derived from the more robust consolidated UMAP (replicates 1–3, 5 and 6). Gene set enrichment analysis per Leiden cluster or treatment was based on the normalized Wilcoxon-Mann-Whitney statistic computed using singleseqgset v0.1.2.9.

To analyze differential proportions of ribosomal RNA expression between treatments, we trimmed original 3’ reads to 75 bp and 5.8S, 18S, and 28S reference sequences on chromosomes 1, 5, 7, 11, and 13. Mapping via HISAT2 (version 2.0.4) allowed no spliced alignment (specified with the argument ‘--no-spliced-alignment’), so that only reads mapping uniquely to genomic loci were retained. Although not polyadenylated, rRNAs are so numerous as to bind non-specifically to beads and can be cautiously analyzed in this manner (21).

## Results

### An additional blood meal increases P. falciparum sporozoite infectivity to hepatocytes

To investigate the effects of an additional blood meal on sporozoite infectivity to liver cells, we provided *An. gambiae* mosquitoes with an infectious blood meal containing *P. falciparum* NF54 gametocytes and then three days after infection, gave an additional (uninfected) blood meal to approximately half of these mosquitoes (2BF group), while the remainder females were maintained on sugar (1BF group) (**Figure 1A**). In total, six experiments were performed: three compared unmanipulated 1BF mosquitoes to ds*GFP*-injected 2BF mosquitoes, which served as controls in a parallel experiment not reported here; and three compared unmanipulated 1BF and 2BF mosquitoes. While infection intensity varied between batches, relationships between ds*GFP*-injected and unmanipulated 2BF groups and their 1BF comparators were identical (**Figure S1**).

**Figure 1.**
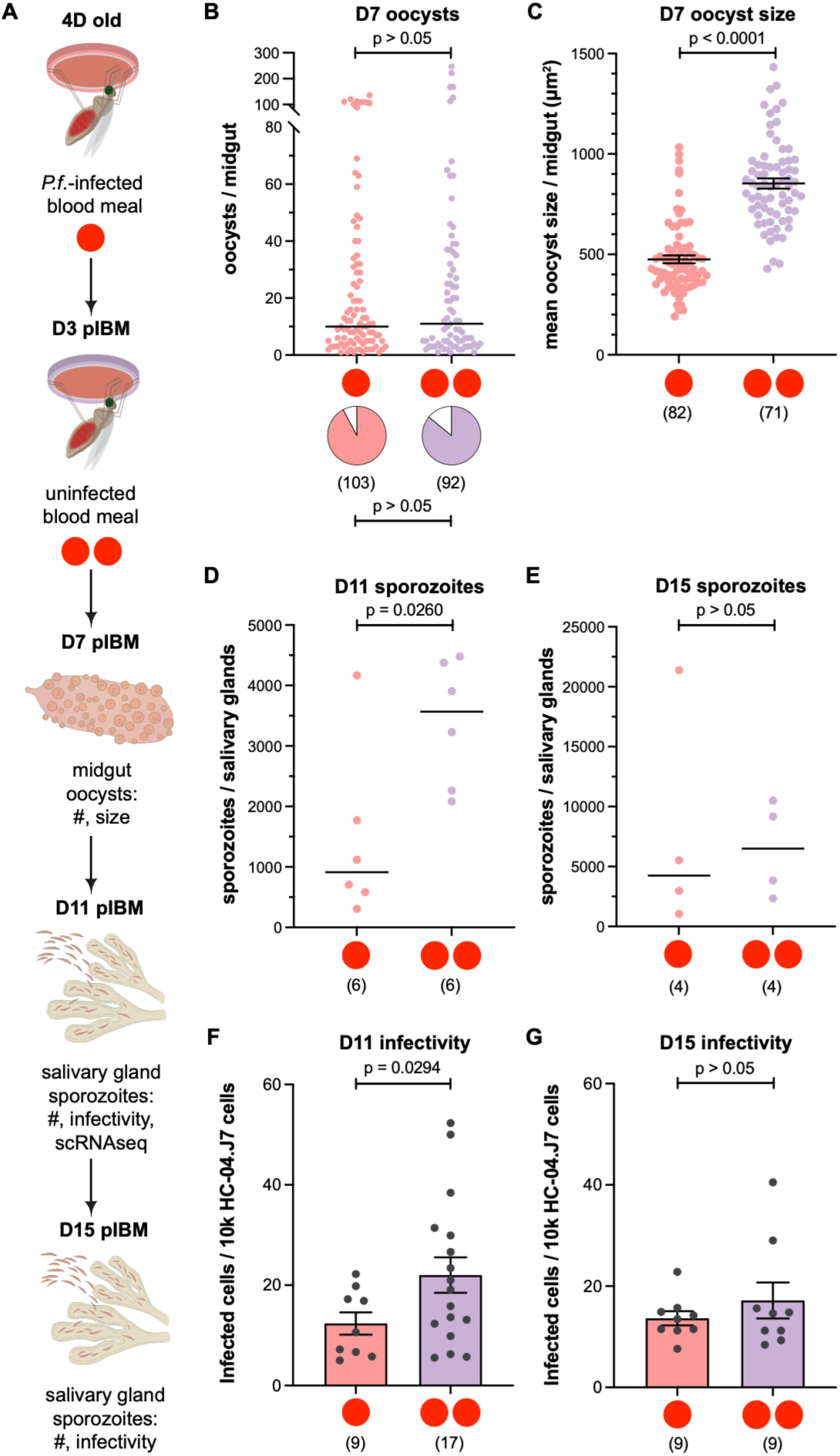
An additional blood meal increases the infectivity of *P. falciparum* sporozoites. (A) Schematic outline of the experimental design. 4-day-old mosquitoes were provided with a *P. falciparum* (*P.f.*) infectious blood meal (single red dot), then on 3 d pIBM approximately half were provided with an additional uninfected blood meal (double red dot), while the remainder were maintained on 10% glucose solution. (B–E) The prevalence (pie-charts, Fisher’s exact) and intensity (Mann-Whitney) of oocysts was unchanged at 7 d (B) but their mean oocyst size was significantly increased (C) (Welch’s t, unequal variances, log-transformed), leading to an increase in the number of salivary gland sporozoites at 11 d (D) (Mann-Whitney), but not by 15 d (E) (Mann-Whitney). Data shown are pooled from 6 infection experiments. Mean oocyst size data (C) excludes midguts with fewer than 3 oocysts. Salivary glands were pooled and purified sporozoite yield (D–E) was normalized to the number of mosquitoes dissected in each experiment. (F–G) Infectivity of sporozoites from females provided an additional blood meal was increased at 11 d (F) (Welch’s t, unequal variances), but not at 15 d (G) (Unpaired t, log-transformed). Sporozoites were limiting in some experiments, so data shown are pooled from 4 (F) or 3 (G) infection experiments. See also **Table S1**.

As previously observed, oocyst size at 7 d pIBM was increased in 2BF females compared to 1BF females, whereas oocyst numbers were unchanged (**Figure 1B–C**). Consistently, at an early timepoint during salivary gland invasion (11 d pIBM), total numbers of sporozoites dissected from salivary glands were increased in 2BF groups, again as expected (**Figure 1D; Table S1**).

We next asked whether sporozoite infectivity is affected by a second blood meal. To this end, we used equal numbers of sporozoites from the two groups to infect HC04.J7 human hepatocarcinoma cells (28), and determined their intra- or extracellular localization using an immunofluorescence assay (30). After 24 h, rates of infection were significantly higher (1.8-fold) in hepatocytes infected with 2BF sporozoites compared to the 1BF control group (**Figure 1F**). Salivary glands dissected at a later time point (15 d pIBM) had similar intensity of infection in the two groups (**Figure 1E**), and sporozoites harvested at this time point showed no difference in infectivity to hepatocytes (**Figure 1G**). All together, these results suggest that additional nutrient availability during parasite development boosts sporozoite infectivity early in infection, an important finding given the high rates of multiple blood feedings in the field (2).

### scRNA-seq analysis identifies multiple sporozoite subpopulations within the salivary glands

To identify the factors that may be responsible for driving differential infectivity after an additional blood meal, we compared single cell transcriptomics (scRNA-seq, 10x Genomics) profiles of sporozoites populations from 1BF and 2BF groups at 11 d pIBM, the time point when we saw increased infectivity in sporozoites derived from 2BF females. We obtained between 573–931 good quality cells (>400 UMI, >125 genes detected, <0.025 proportion mitochondrial transcripts) in each biological replicate (except replicate 4, where no scRNA-seq was performed) (**Table S1**). After random subsampling to equalize cell counts between the two groups per replicate (average = 616), the dataset averaged 197 genes and 541 UMIs per sporozoite. We also generated a single 15 d pIBM comparator sample featuring similar scRNA-seq metrics (**Table S1**).

As an initial step, we analyzed all the available data at 11 d pIBM as a whole (pooling data from 1BF and 2BF mosquitoes), which identified a substantial degree of variation in gene expression between individual sporozoites (**Figure 2A**). This analysis revealed a gradient of gene expression that recapitulated all stages of mosquito-stage sporozoite development previously described ranging from day 12 midgut sporozoites (ooSpz) to day 20 salivary gland sporozoites incubated in fibroblast growth media (actSpz) (22). After RPCA integration, we identified different clusters of salivary gland sporozoites; a smaller cluster (labelled 1) and a second much larger cluster containing more extensive variation, which we therefore split further into two sub-clusters (labelled 2 and 3) (**Figure 2A**). Using pseudotime analysis (31), we ordered gene expression profiles and described a transcriptional trajectory from cluster 1 to 3, anchoring the pseudotime axis based on sequencing derived from the sporozoites harvested at 15 d pIBM (**Figure 2B; Figure S4**). While a single experiment, the day 15 data were consistent with the directionality we inferred below from marker genes within our day 11 samples. We identified 61 genes whose expression strongly correlated with this transcriptional trajectory (**Figure S5**), including early genes associated with sporozoite egress from oocysts and invasion of the salivary glands (e.g. MAEBL (32, 33), TRP1 (34), CRMP2(35) and CRMP4 (36)), while transcripts expressed later were associated with development in hepatocytes (e.g. LSAP2 (37), PTEX150 (38) and EXP1 (39)), suggesting parasites increase expression of genes in anticipation of their subsequent requirement in the liver. Note that UMAP and pseudotime plots using sporozoite data from only unmanipulated mosquitoes were equivalent to **Figure 2**, with a smaller immature cluster and a larger more mature cluster, indicating no ds*GFP*-related (**Figure S3**) or oocyst intensity-related (**Figure S1**) effects.

**Figure 2:**
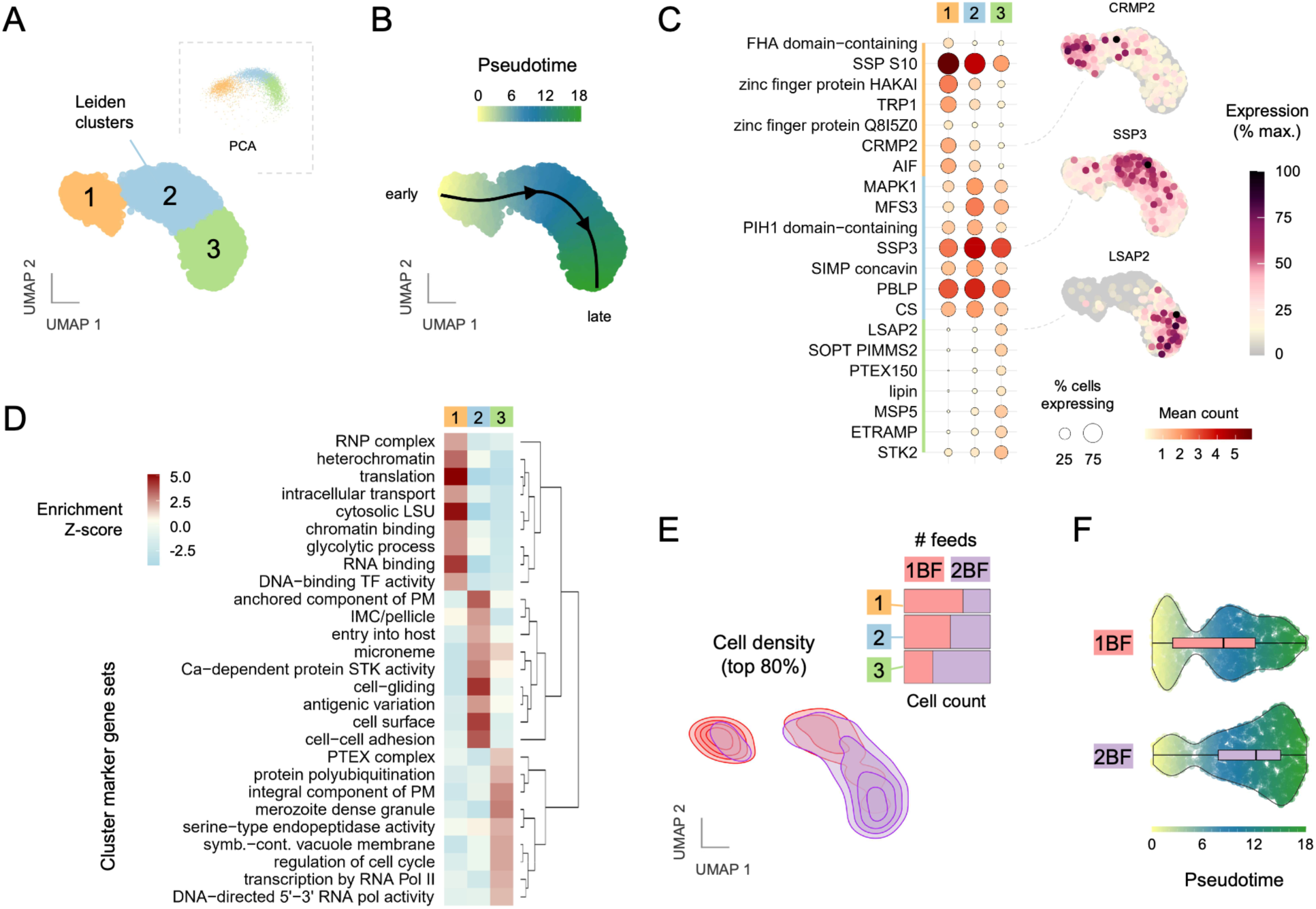
***P. falciparum* salivary gland sporozoites show a developmental transcriptional trajectory.** 11-day sporozoite transcriptional profiles (pooled from 5 experiments) form 3 clusters by UMAP projection (A: inset, Principal Component Analysis (PCA) plot) that are connected over a single developmental pseudotime trajectory (B, arbitrary units). (C) Mean read counts for the top 7 marker genes with annotated functions and an average log_2_FC > 0.585 in expression over other clusters (across all samples) and p < 0.001 are shown for the 3 clusters. Percentage of maximal expression is shown overlaid on the UMAP for an example gene from each cluster. Percentage of cells expressing each gene is shown by dot size. (D) GO term enrichment and clustering analysis showing GO terms of gene sets (>5 genes) with an average log_2_FC > 0.585 over expression in other clusters. (E) Kernal density UMAP showing an additional blood meal shifts cellular occupancy to later clusters with more mature transcriptional profiles. Most dense 80% of cells plotted. Inset, mosaic plot showing cell counts differ significantly by cluster and blood feed treatment (chi-squared statistic χ2 = 193.6, d.f. = 2, p < 0.0001) (F) Violin plots showing differences in cell abundance across pseudotime (early, left to late, right). Internal box plot shows the median and interquartile range.

Many of the pseudotime-associated genes overlapped with those identified through our analysis of marker genes of each cluster (**Figure 2C; Figure S5; Table S3**). Again, we found that genes expressed earlier (cluster 1) included those associated with salivary gland invasion (e.g. CRMP2, TRP1) whereas genes expressed in cluster 3 were associated with more mature sporozoites and liver-stage development (e.g. LSAP2, PTEX150). Clusters overlapped with all previously identified groups of sporozoites (22), with cluster 1 overlapping most closely with oocyst sporozoites (ooSpz) and clusters 2 and 3 overlapping with hemolymph (hlSpz) and salivary gland (sgSpz) sporozoites (**Figure S6**). GO term enrichment analysis of each subcluster (**Figure 2D; Table S4**) revealed the early importance of the regulation of gene expression at heterochromatic and translation levels, as sporozoites establish a translationally repressed state in the salivary glands. Cluster 2 was enriched in GO terms associated with cell motility, as well as with cell surface proteins and cell invasion processes, as sporozoites change their surface proteins in anticipation of host entry (**Figure 2D, middle column**). The most mature cluster 3 was labeled with processes fitting with transcription, replication and protein export in the liver stage, as sporozoites prepare to establish liver stages (**Figure 2D, right column**).

### An additional blood meal shifts salivary gland sporozoites to later transcriptional profiles

We next compared how an additional blood meal affects the abundance of cells within each cluster and their transcriptional profiles. To be confident of identifying blood-feeding induced changes in sporozoite gene expression relevant to transmission settings, we specifically focused our analysis on replicates 5 and 6, in which sporozoites were harvested from unmanipulated mosquitoes with field-like infection intensities fed once or twice (**Figure S1**). Nevertheless, differentially expressed genes and fold changes identified using data pooled from all replicates were highly similar (**Figure S3C; Table S2**), suggesting limited density-dependent impacts of higher oocyst intensities on sporozoite transcription. Although all clusters were present in sporozoites from mosquitoes fed once or twice, an additional blood meal significantly shifted the cellular occupancy such that a greater proportion of sporozoites in the 2BF group had a late cluster 3 transcriptional profile (chi-squared statistic χ2 = 193.6, d.f. = 2, p < 0.0001; **Figure 2E–F**). Sporozoites from the 1BF group were largely found in the earlier cluster 1, whereas cells from the 2BF group were mostly in cluster 3, shifting the median pseudotime value (**Figure 2F**). Therefore, the additional resources acquired from a second blood meal not only increase numbers of sporozoites found within the salivary glands at an early timepoint (3), but also promote their transcriptional readiness for their next developmental transition, in turn increasing their infectivity.

### Transcripts differentially regulated by an additional blood meal relate to cell motility, surface protein remodeling and liver-stage development

We went on to analyze differences in sporozoite transcriptional profiles between blood feeding regimens using ScranFindMarkers (ScranFM), which controls for batch differences between experimental replicates. We performed analysis at two levels: globally, pooling cells across all clusters; and within-clusters, directly comparing cells at similar developmental states (described later). While fold changes were modest between treatment groups (cut-off at 1.2-fold and a highly stringent adjusted p-value < 0.0005), the global analysis revealed 131 genes significantly upregulated and 45 genes significantly downregulated in sporozoites from the 2BF group (**Figure 3; Table S5**), with 34 (26%) and 13 (29%), respectively, annotated with unknown functions. Many of these differentially expressed genes are likely relevant to sporozoite functions, as they show either limited expression during earlier mosquito stages, or turn on in sporulating oocysts (**Figure S7**) (40).

**Figure 3:**
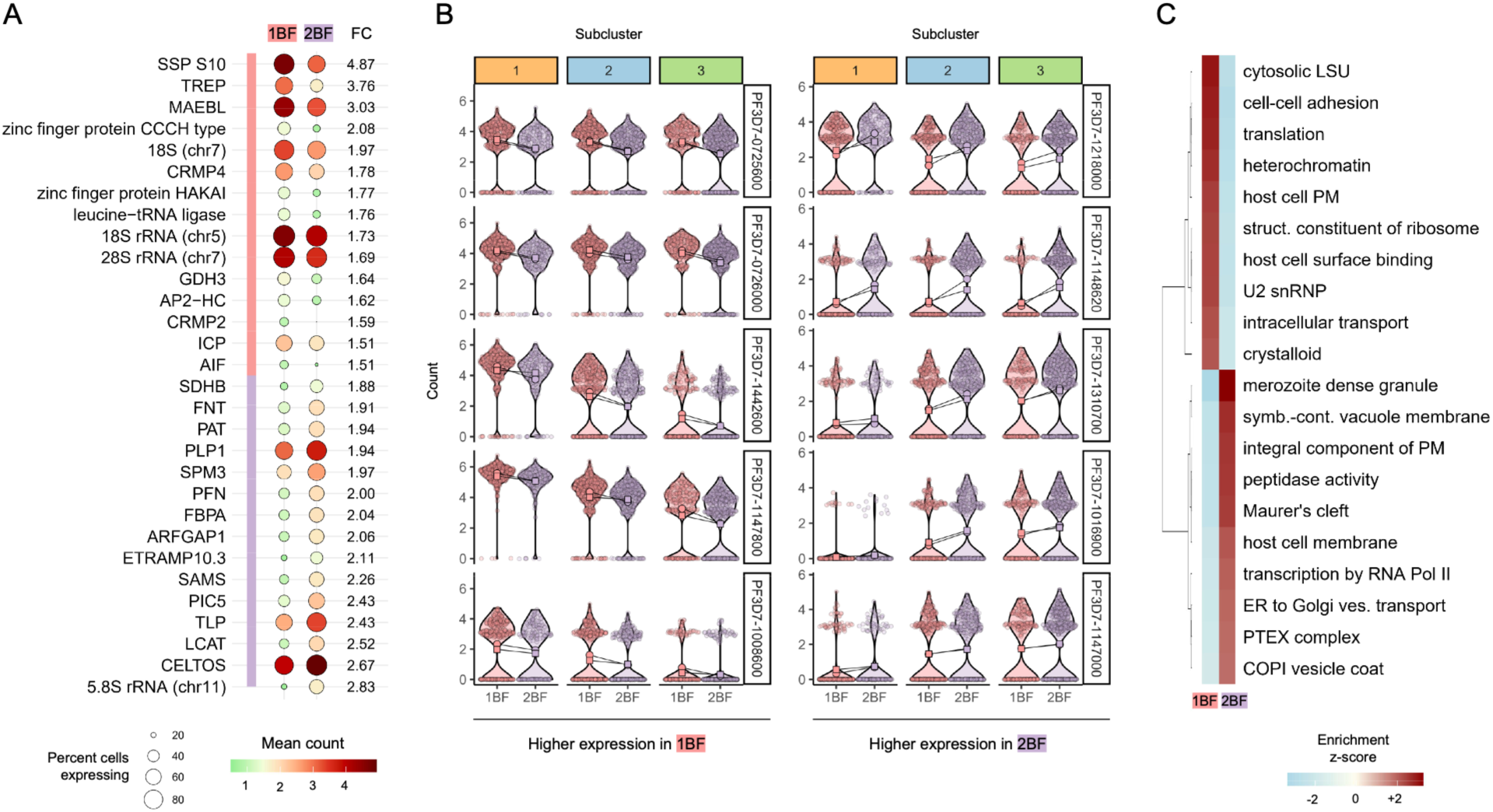
An additional blood meal shifts *P. falciparum* sporozoites to a more mature transcriptional profile. (A) Mean read counts are shown for 15 selected up- and down-regulated genes with annotated functions, a p-value < 0.0005, and a FC > 1.5 in expression when comparing between 1BF and 2BF treatments (global analysis across all clusters). Percentage of cells expressing each gene is shown by dot size. (B) Violin plots showing read counts of differentially expressed genes in sporozoites across clusters (1, early to 3, late). Lines connect median expression values between 1BF and 2BF treatments (circle: mean, replicate 5; square: mean, replicate 6). (C) GO term enrichment and clustering analysis showing GO terms of gene sets (>5 genes) with an average log_2_FC > 0.585 when comparing between 1BF and 2BF treatments (global analysis across all clusters).

In 2BF sporozoites we detected a strong signature of upregulation in genes associated with cell motility, including: FBPA, a glycolytic enzyme and known binding partner of TRAP in the sporozoite gliding motility complex, known as the glideosome (41); Actin (ACT1), the cytoskeletal protein subunit; Profilin, an actin-binding protein required for motile force generation (42); and PAT, a regulator of surface ligand exocytosis important for secreting TRAP, and without which sporozoites are immotile (43) (**Figure 3A**). Components of the inner membrane complex (IMC), a membranous organelle lying beneath the plasma membrane and supported by the cytoskeleton (44), were also upregulated in the 2BF group: SPM3, which in *P. berghei* has been shown to be required for the association of the IMC with the sub-pellicular microtubule network and may be involved in generating motile force (45); PIC5 (PF3D7_1310700), a component of the PhIL1-interacting protein complex in the glideosome, which has a role in merozoite motility that here may extend to sporozoites (**Figure 3A**; **Figure 3B, right panel**)(46). We also identified a non-coding RNA, PF3D7_0108900, antisense to a 600 bp region encompassing the first exon of PhIL1, that could potentially regulate expression of this motility complex. Together, these upregulated genes suggest transcription of cellular components is increased to convey a greater ability to generate cell motility in 2BF sporozoites.

Many well-annotated surface ligands and protein families were also increased in expression in sporozoites from 2BF mosquitoes: TRAP-like protein (TLP) (47), CelTOS (48), SPATR (49), AMA1 (50), MSP4 (51), and MSP5 (52) (**Figure 3A**). Several of these surface ligands require proteolytic cleavage for their functionality and the upregulated proteases ROM1 (53) and subtilisin 2 (SUB2) (54) could play a role in activating these or other surface ligands to promote infectivity. Interestingly, we also found a known substrate of SUB2, PTRAMP (PF3D7_1218000) (55) (**Figure3B, right panel**) and its interacting partner, CSS (56, 57), suggesting these micronemal proteins could be important for sporozoite as well as merozoite invasion processes. Conceivably, the increased expression of functional surface ligands may contribute to the enhanced sporozoite infectivity we observed.

As well as components for liver cell invasion, 2BF sporozoites also showed upregulation of genes involved in liver-stage development. SLARP (PF3D7_1147000), a master regulator of merozoite development in hepatocytes (58) was increased (**Figure 3B, right panel**). Several other exported proteins localized to the parasitophorous vacuole (PV) and its membrane during this stage were also identified in the upregulated gene set, such as PV1(59), PV3(60), and ETRAMP10.3 (PF3D7_1016900)(61) (**Figure 3B, right panel**), along with components of the cellular protein ER translocation and export machinery: PTEX150, a known interacting partner of PV1(59), and co-chaperone interactors SEC63 (62) and Pfj2 (63, 64). GO term enrichment analysis of the upregulated 2BF genes also showed a strong signal for vesicle trafficking and integral proteins within cellular and host membranes (**Figure 3C, Table S6**). Together, the production of these transcripts in transmission stages could increase the speed of functional assembly of transport complexes in hepatocytes and increase the likelihood of parasite survival.

Among the genes downregulated in 2BF females were several with known roles in salivary gland invasion (**Figure 3A**; **Figure 3B, left panel**), such as MAEBL (PF3D7_1147800) (32, 33), sporozoite specific protein S6, also known as TREP (PF3D7_1442600) (65), CRMP2 (35) and CRMP4 (36). Their decreased expression makes biological sense given these genes are no longer required following salivary gland invasion. More generally, transcripts associated with the gene expression machinery, such as 18S and 28S rRNAs (with the exception of 5.8S (PF3D7_1148620), leucine-tRNA ligase, and the pre-rRNA processing factor PF3D7_0931700 were all more lowly expressed in the 2BF group as compared to the 1BF group, consistent with the previously reported decrease in translational activity in more mature sporozoites (10–13, 18, 19) (**Figure 3B, left panel**). GO term enrichment analysis also pointed to reduced gene expression, with terms for ‘heterochromatin’, ‘transcription by RNA PolII’ and ‘cytosolic large [ribosomal] subunit’ all enriched (**Figure 3C; Table S6**). Ribosomal RNAs are transcribed from 5 genomic loci in *P. falciparum*, distinguished by their stage-predominant expression patterns: asexual stages express from A-type loci (chromosomes 5, 7), whereas sexual stages express from the S1 locus (chromosome 1) during gametocyte to oocyst stages and transition to the S2 loci (chromosomes 11, 13) in oocysts and sporozoites (66). The 28S rRNA locus on chromosome 8 had a negligible expression in our dataset. While not specifically targeted by the poly(dT) UMI oligos on 10x gel beads, these transcripts are abundant in scRNA-seq datasets (21). We analyzed the proportion of rRNA reads mapping unambiguously to the above-described rRNA loci and consistently observed a shift away from transcription of A-type rRNAs and towards S-type rRNAs in sporozoites from the 2BF group (**Figure 3B; Figure S8A**). This evidence supports the hypothesis that these sporozoites are more transcriptionally mature than sporozoites derived from mosquitoes fed once, with the caveat that such an inference from raw reads without UMI counts requires caution. Interestingly, although transcribed as polycistronic precursors (in the order 18S-5.8S-28S rRNA) that are later processed, we detected strong differences in the origin of rRNAs between the constituent genes, with the highest proportions of 18S and 5.8S rRNA coming from the S2 locus, while 28S rRNA from the S1 locus predominated (**Figure S8**), potentially reflecting a transition from S1 to S2 locus transcription.

### Cluster-specific analysis validates bloodmeal-driven gene expression changes beyond pseudotime distribution effects

To verify our differentially expressed genes were not solely a function of differential sporozoite abundance between pseudotime extremes, we repeated the differential gene expression analysis within each cluster (c1, c2 or c3) using ScranFM, comparing sporozoites at similar developmental stages between the 1BF or 2BF treatment groups. Owing to fewer cells in each cluster, we used a more relaxed p-value threshold (p<0.05) and identified slightly more differentially expressed genes (**Figure S9; Table S7**). Although some previously identified genes of interest (e.g. SLARP, PTEX150) were no longer identified, 65% (127/196) overlapped with our global analysis, importantly confirming that changes to sporozoite transcripts between blood feeding regimens persist when considering similar developmental states, rather than reflecting unequal sporozoite distribution across clusters. Genes shared across multiple clusters had consistent changes to their expression over pseudotime, and were often previously identified in the global analysis (85%, 55/65); these encoded more basic cellular functions (e.g. ribosomal rRNAs, heat shock proteins, eIF5, histone 2B; **Figure 3B, top 2 rows**). Other genes showed steady increases or decreases across pseudotime (**Figure 3B, bottom 3 rows**), suggesting that an additional blood meal may first affect more fundamental cellular processes driving oocyst growth, that carry over into the transcript levels detected within all sporozoite clusters, with more cluster-specific changes reflecting potentially more infectivity-relevant changes.

This cluster-focused approach also identified genes not present in our global analysis, either because bloodmeal-induced changes were too small to initially be detected, or their regulation is stage-specific rather than temporal. In cluster 1, transcripts for oocyst rupture protein 2 (ORP2) were more highly expressed in 2BF sporozoites, consistent with its role in sporozoite egress from the oocyst (67). In cluster 2, several regulators of gene expression were elevated after a second blood meal, suggesting that transcript repression is stronger: eIK2, a kinase responsible for translation repression in sporozoites (10); an RNA-binding protein of infectious sporozoites, UIS12, required to stabilize stored transcripts (16, 68); HMGB2, a small DNA binding protein that helps activate or repress gene expression (69), and the histone H3 variant H3.3, which marks the promoter regions of inactive but poised genes (70). We also identified an increase in HoMu, an RNA-binding protein associated with mRNAs stored in ribonucleoprotein complexes (71) and HMGB1 in cluster 3.

In the opposing direction, surprisingly, several genes linked to sporozoite function and/or transmission were lower in the more mature clusters 2 and 3. These included: SERA8 a cysteine protease required for oocyst egress and establishment of liver infection (72); GEST, a secreted protein required for hepatocyte traversal (73); and SPELD, a surface protein required for EEF growth (74). Despite established roles in sporozoite function and/or liver-stage development, their reduced expression in 2BF sporozoites may reflect accelerated transcript degradation in advance of protein function, likely following sufficient protein accumulation on the sporozoite surface or in secretory bodies (73, 74).

## Discussion

In this study we find that an additional non-infectious blood meal accelerates the acquisition of infectivity of *P. falciparum* salivary gland sporozoites by shifting their transcriptional profile to one consistent with mature sporozoites. This effect may be due to a combination of factors. Firstly, by accelerating oocyst development, sporozoite segmentation and maturation can occur in a shorter timeframe, and a greater number and proportion of cells present within the salivary glands will show enhanced infectivity and its underlying transcriptional profiles. Secondly, additional resources during oocyst development may trigger higher expression of key transcripts required for infectivity. We discuss these two main explanations considering the literature below.

The invasion of the salivary glands has been found to be a key step in the development of maximal infectivity. Previous work examining sporozoite maturation has found lower levels of motility and infectivity in immature sporozoites from midgut oocysts and the hemolymph compared to salivary gland populations (5, 6, 15, 75, 76). Two lines of evidence had suggested that maturation within the salivary glands *per se* was not required for infectivity in *P. gallinaceum and P. berghei* model species, as both *in vitro*-produced sporozoites and hemolymph sporozoites (14 and 17–22 d pIBM) were infectious by intravenous injection (4, 76–78). Additionally, motile *P. berghei* mutants unable to invade the salivary glands (*maebl^−^*, *crmp1^−^* and *crmp2^−^*mutants) remained infectious to hepatoma cells or rodents by injection, further suggesting that salivary gland invasion was not required for transmission (32, 35). However*, P. berghei* hemolymph sporozoites are defective in motility (75, 76), and functional comparisons in hepatocyte traversal and *in vivo* infection show oocyst and hemolymph sporozoites have limited infectivity, and that gland invasion is required for optimal transmission (15). Similar results were found comparing oocyst and salivary gland sporozoites in *P. gallinace*um infections of chickens (5). Furthermore, in *P. falciparum*, hemolymph sporozoites sampled over several days did not improve their hepatocyte traversal rates (79), suggesting they lacked a key trigger of full maturation. By accelerating oocyst development, an additional blood meal will allow invasion and maturation within the salivary glands sooner. In future studies, it will be interesting to identify the molecular events during invasion triggering the transcriptional/translational changes of full maturation.

Evidence from isolated studies suggests time spent within the salivary glands impacts infectivity, especially in the first days. Previous work has observed a gradual increase in *P. berghei* sporozoite infectivity with time in the salivary glands (4), and several days’ incubation therein is required for optimal transmission to mice (15). By bringing gland invasion forward in time, an additional blood meal will extend the total time spent within the salivary glands. Interestingly, additional blood feeding events have been deliberately incorporated to boost sporozoite yields in laboratory settings (80). Faster-developing *P. falciparum* sporozoites in females with impaired reproduction remain infectious to primary human hepatocytes (81), while sporozoites kept within the salivary glands beyond 14 days (82) or over several weeks (83) decline in infectivity, in contrast to *P. yoelii* (84) or *P. berghei* (4, 15). Additional feedings could potentially preserve sporozoite infectivity over a more extended period in *P. falciparum*, by, for example, reinforcing gene regulation by translational repression through eIK2 (10).

Infectivity of sporozoites has also been correlated to the parasite load in salivary glands (85, 86), with a large increase in transmission observed in some studies once numbers surpass ∼10,000 parasites (87, 88). This was attributed to an increase in the sporozoite number in each inoculum (85–88). An additional blood meal will result in earlier crossing of such a threshold as it increases sporozoite loads present by a given time (3, 89–93). However, as we standardized sporozoite numbers in our hepatocyte infectivity assay, our data suggest that increased infectivity *per sporozoite* can also contribute to increased transmission by multiply-fed females, at least early in infection, as the higher sporozoite loads would also be associated with a greater proportion of mature sporozoites.

While several studies have approached either sporozoite infectivity or gene expression in the context of an additional blood meal, none has addressed this question as directly as we have in *P. falciparum* through our simultaneous infectivity assays and scRNA-seq studies. Early work found a boost to hepatocyte invasion of *P. falciparum* sporozoites following an additional blood meal 5 d pIBM when it contained anti-*Pf* antibodies, over unrelated antibodies, as this stimulated higher oocyst productivity (sporozoites/oocyst) and salivary gland loads (94). However, changes to oocyst productivity were not observed by others in similar experiments (10 d pIBM (95); 5–11 d pIBM (89)), and no 1BF treatment group was provided as a comparator for infectivity (94). Costa et al., (96) have shown an additional blood meal at 7 d pIBM rescues the effects that depletion of the mosquito lipid transporter Lipophorin has on *P. berghei* oocyst size and sporozoite load and infectivity to mice. In their work, comparison between 1BF and 2BF control treatments at day 18 showed no increase in EEF numbers in 2BF sporozoites (96), but besides using a different parasite species, this is likely because by this timepoint both treatment groups have reached full maturity, as we see in *P. falciparum* in our day 15 dataset. Our data confirm that sporozoites undergo a developmental gradient of maturation in the salivary glands, from transcriptional profiles consistent with premature, recently-invaded sporozoites through to mature sporozoites upregulating critical liver-stage genes (**Figure 2**). Previous scRNA-seq data at later timepoints post infection (*P.f.*: > 12–20 d pIBM; *P.b.*: 14–26 d pIBM; *P.v.*: 16–18 d pIBM) also identified a series of transcriptional states within sporozoites (20–23). Even at the early day 11 timepoint, we find three clusters representing all stages of sporozoite maturation within the salivary glands, indicating transcriptionally premature sporozoites characteristic of the oocyst are capable of salivary gland invasion and no lengthy maturation in the hemolymph may be necessary (**Figure S6**). An additional blood meal shifts sporozoites forward along this maturation gradient. Immature transcriptional profiles may represent recently invaded sporozoites, perhaps localized under the salivary gland basal lamina prior to invading acinar cells. RON11^−^ mutant parasites deficient in salivary gland invasion have been found to accumulate in this initial compartment (97). Alternatively, parasites may be closely associated or attached to the salivary gland surface and in the process of invasion. We consider it unlikely that these immature sporozoites represent contaminating hemolymph sporozoites as they are less abundant in 2BF mosquitoes, in which accelerated oocyst development would be expected to have significantly increased their numbers at this early time point in sporozoite invasion of salivary glands. Additionally, our data are consistent with a study examining *P. vivax* salivary gland sporozoites (day 16–18), which identified two large clusters of sporozoites, one more immature than the other (23) (we further subclassified the larger cluster into 2 sub-clusters).

We identify over a hundred transcripts, many of which encode genes of unknown function, significantly changing in 2BF sporozoites that may mediate this boost in infectivity, largely through accelerated maturation (**Figure 3**). Highly overlapping lists of genes were recovered in analyses both at a broad resolution across day 11 salivary gland populations as a whole, and within each identified cluster, suggesting they are not due to differential abundance of mature sporozoites between treatments, although we cannot control for small differences in maturation with only 3 Leiden clusters (**Figure S9**). While we are unable to confidently assert that 2BF sporozoites are qualitatively different from 1BF sporozoites at day 11, our cluster-specific analysis may contain some bloodmeal-induced changes to the transcriptional route to infectivity that we are not able to fully resolve from temporal effects (small differences in sporozoite maturity or salivary gland residency). The persistence of any transcriptional differences also remains unknown as sporozoites from both 1BF and 2BF groups mature: infectivity is no longer different between blood feeding regimens at day 15 (**Figure 1G**), and we were regrettably unable to generate additional sequencing libraries at this timepoint. Whether an additional bloodmeal produces qualitatively different sporozoites therefore remains an outstanding question.

We identified transcripts associated with cell motility, surface proteins, gene expression regulation, and liver-stage development as upregulated by an additional blood meal. Of note, transcripts of PTRAMP (55) and its interacting partner CSS (57), members of the PCRCR merozoite invasion complex that binds basigin on the surface of human erythrocytes were upregulated (56, 57, 98). While these transcripts may be being produced well in advance of their requirement in developing merozoites, the expression we detect in sporozoites suggests a potential role in other invasive steps with an unknown ligand (18) that has not previously been investigated. Upregulation of other transcripts of genes known to be localized to the parasitophorous vacuole in the liver stage (and therefore unlikely to be translated in sporozoites), and of the translational repressor eIK2 (10), suggests parasites store a greater number of transcripts after an additional blood meal to prepare for the subsequent stage of parasite development. Oocysts sensing nutrient availability adjust gene expression machinery (92) to accelerate growth and increase transcript numbers of key genes required for infectivity prior to sporozoite individualization. Indeed, higher expression of a fluorescent maturation reporter in sporozoites has been correlated to greater infectivity in salivated sporozoites and a larger size of the resultant EEFs (15). Infectivity has also been linked to sporozoite mitochondrial function in *P. berghei* (96), but not in *P. falciparum* (82) and we did not detect a significant enrichment of genes associated with energy production in our data, suggesting this is not regulated, at least at the transcriptional level, in this species.

We observed minimal differences in oocyst size and bloodmeal-induced changes to the sporozoite transcriptome between low intensity infections and our pooled data incorporating high intensity infections (**Figure S1; Figure S3**). This suggests density-dependent effects do not negatively impact *P. falciparum* oocyst development (81), unlike *P. berghei* oocysts (92), to potentially limit sporozoite infectivity. Future work could also explore the impact of additional feedings on sporozoites already resident in the salivary glands and any density-dependent effects in this tissue.

We believe our results are highly relevant for understanding transmission in natural populations, where female lifespan is short and multiple blood feeding occurs frequently (2). Not only are sporozoites numbers increased in quantity in the salivary glands at earlier timepoints following an additional blood meal (3), but their infectivity (quality) is also increased, further enhancing parasite fitness and chances of transmission. This suggests parasites are rapidly able to adjust their development to exploit resources when available to improve their odds of transmission, a finding that confirms previous work demonstrating that *P. falciparum* parasites are incredibly well adapted to their *Anopheles* mosquito vectors (81).

## Acknowledgements

We would like to thank Elizabeth Nelson, Kaileigh Bumpus, and Aaron Stanton for *An. gambiae* mosquito rearing and Naresh Singh for *P. falciparum* gametocyte culturing. We would also like to thank other members of the Neafsey and Catteruccia laboratories, and 10x Genomics for helpful discussions on this project. We also thank Rhoel R. Dinglasan for the HC04-J7 human hepatocarcinoma cell line. The monoclonal α-*Pf*CSP, clone 2A10 (MRA-183A), previously contributed by Elizabeth Nardin, was obtained through BEI Resources, NIAID, NIH.

## Author Contributions

Conceptualization: W.R.S., P.S., D.E.N., F.C.;

Data Curation: W.R.S., P.S.;

Methodology: W.R.S., P.S., D.E.N., F.C.;

Resources:

Investigation: W.R.S., P.S., M.A.I., S.A., J.K.;

Formal analysis: W.R.S., P.S.; L.H.V., Y.Y.;

Software:

Validation:

Writing – Original Draft: W.R.S., F.C.;

Writing – Review and Editing: W.R.S., P.S., D.E.N., F.C.;

Visualization: W.R.S., P.S.;

Supervision: D.E.N., F.C.;

Funding acquisition: D.E.N., F.C..

## Declaration of Competing Interests

The authors declare that no competing interests exist.

## Data Availability

Numerical data supporting mosquito and hepatocyte infection assays is available on Harvard Dataverse at X. scRNA-seq data is publicly available at SRA16272983. Code used to analyze the data is available at https://github.com/fishntryps/spz_scRNAseq.

## Funding Information

This study was supported with federal funds from the National Institute of Allergy and Infectious Diseases, National Institutes of Health, Department of Health and Human Services, under grant numbers U19AI110818 to the Broad Institute and managed by D.E.N. and R01AI148646 and R01AI153404 to F.C.. F.C. is funded by the Howard Hughes Medical Institute (HHMI, www.hhmi.org) as an HHMI Investigator. Y.Y. is funded by the Charles A. King Trust Postdoctoral Research Fellowship Program, Bank of America Private Trust, Trustee. The findings and conclusions within this publication are those of the authors and do not necessarily reflect positions or policies of the HHMI, the NIH or the Bank of America Private Bank. The funders had no role in the study design, in data collection, analysis or interpretation, in the decision to publish, or the preparation of the manuscript.

## Supplementary Table Legends

**Table S1: Details of sample collection, hepatocyte infection assays and scRNA-seq.** All samples were dissected salivary glands from *Anopheles gambiae* G3 females infected with *Plasmodium falciparum* NF54 strain. Sporozoites were occasionally insufficient, or debris was too abundant for adequate single-cell cDNA library construction. †Reads were mapped to the *P. falciparum* 3D7 strain transcriptome (release 58). # = number; BF = blood feed; dsGFP = double-stranded RNA targeting green fluorescent protein; mosq. = mosquitoes; pIBM = post infectious blood meal; spz = sporozoites; UMIs = unique molecular identifiers.

**Table S2: Differentially expressed genes identified using a global scranFM analysis on all replicates (1– 3, 5 and 6).** The table includes all genes identified as differentially expressed in day 11 sporozoites between 1BF and 2BF bloodfeeding regimens with a p-value below 0.0005. Row order descends based on fold-change, which has no threshold cutoff.

**Table S3: Expanded list of cluster-specific marker genes**

The table includes all genes identified as cluster markers with >1.5 fold-change (FC) for member vs. non-member cells (using average FC across the study’s 10 sample collections) and for which p-values fell below 0.05 for all individual samples based on Wilcoxon rank-sum testing with FindConservedMarkers in Seurat v5.0.3. Row order ascends based on maximum p-value (i.e., the weakest p-value observed for any individual sample), grouped by Leiden cluster association (left column). Significance level “***” denotes p < 0.00001, significance level “**” denotes p < 0.001, and significance level “*” denotes p < 0.05. The final column indicates whether the gene is included in the figures visualizing pseudotemporal expression patterning in the 10x data (**Figure 2C**, **Figure 3B**, and **Figure S3**).

**Table S4: Gene set enrichment with respect to Leiden clusters (expanded list)**

The table includes the top 25 gene sets showing enrichment signal per Leiden cluster based on the normalized Wilcoxon-Mann-Whitney statistic computed using singleseqgset v0.1.2.9. Rows are ordered by descending enrichment Z-score, grouped by cluster. For each row (gene set), the p-value associated with the top Z-score is shown, and it is indicated whether the gene set’s cluster-associated expression signal has been visualized in **Figure 2D**. Significance shows “*” if p < 0.05. Note that the table also includes enrichment trends (“.” denotes p > 0.05), primarily in the case of cluster 3.

**Table S5: Differentially expressed genes identified using a global scranFM analysis on replicates with mosquitoes infected with field-like intensities of parasites (5 and 6).** The table includes all genes identified as differentially expressed in day 11 sporozoites between 1BF and 2BF bloodfeeding regimens with a p-value below 0.0005. Row order descends based on fold-change, which has a threshold of 1.2.

**Table S6: Gene set enrichment with respect to feeding condition (expanded list)**

The table includes the top 25 gene sets showing enrichment signal per treatment (# feeds) group based on the normalized Wilcoxon–Mann–Whitney statistic computed using singleseqgset v0.1.2.9. Rows are ordered by descending enrichment Z-score, grouped by treatment. For each row (gene set), the p-value associated with the top Z-score is shown, and it is indicated whether the gene set’s treatment-associated expression signal has been visualized in **Figure 3C**. Significance shows “*” if p < 0.05. Note that the table also includes enrichment trends (“.” denotes p > 0.05) for both treatment groups.

**Table S7: Differentially expressed genes identified using a scranFM analysis on Leiden clusters using replicates with mosquitoes infected with field-like intensities of parasites (5 and 6).** The table includes all genes identified as differentially expressed in day 11 sporozoites between 1BF and 2BF bloodfeeding regimens with a p-value below 0.05, relaxed due to lower numbers of cells. Row order descends based on fold-change, which has a threshold of 1.2. Cluster and pseudotime annotations for cells used in the replicate 5 and 6-focused analysis are based on the consolidated UMAP (replicates 1–3,5 and 6)

## Supplementary Figures

**Figure S1.**
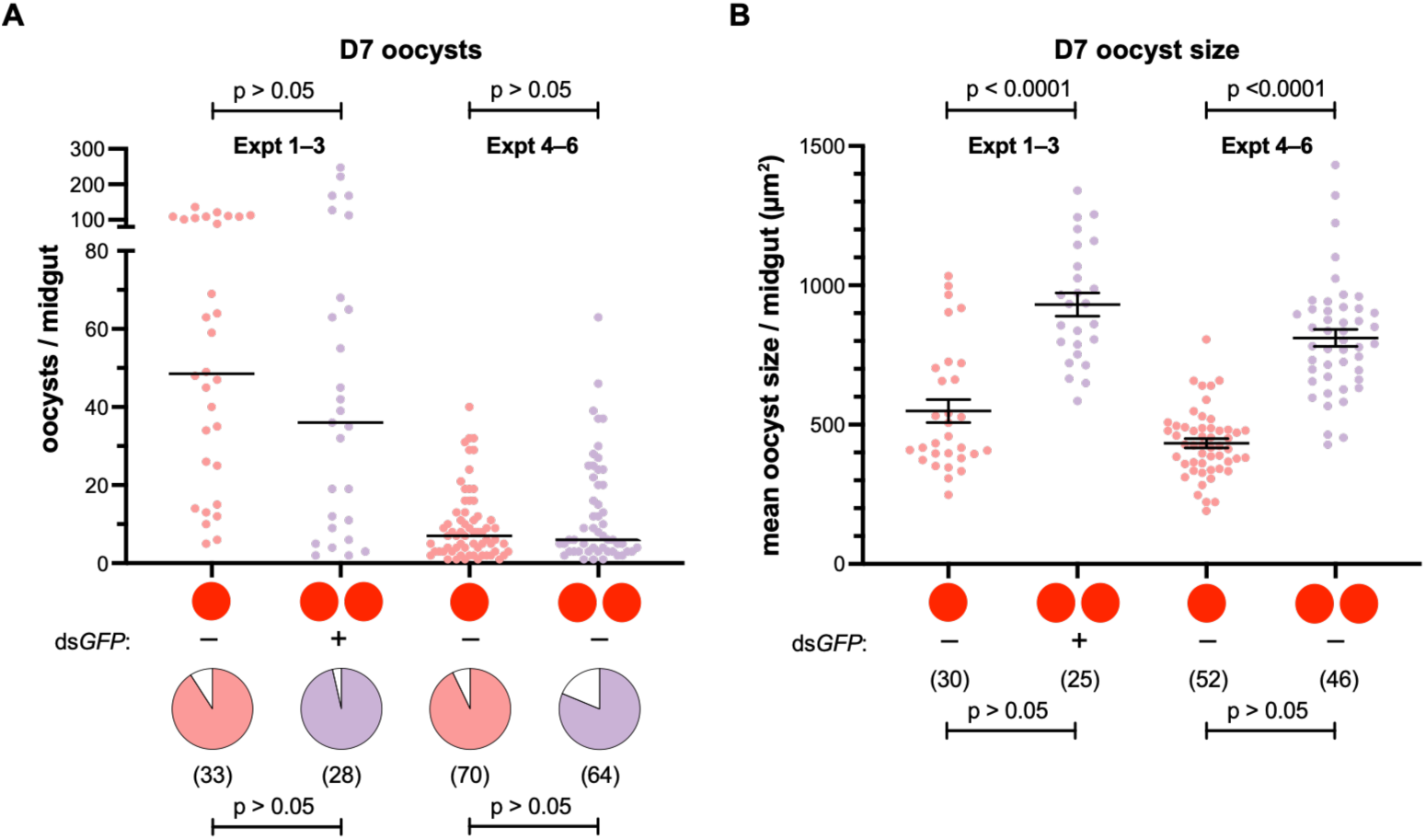
Effects of an additional blood meal at 3 d pIBM on oocyst intensity and size were consistent across experimental batches. The prevalence (Fisher’s exact) and intensity (Mann-Whitney) of oocysts was unchanged at D7 between blood-feeding regimens (A), but mean oocyst size was significantly increased (B) (Welch’s t, unequal variances, log-transformed). Data from each batch are pooled from 3 infection experiments. Experiments (Expt) 1–3 incorporated 2BF ds*GFP*-injected mosquitoes, whereas Expts 4–6 incorporated unmanipulated 2BF mosquitoes. Mean oocyst size data (B) excludes midguts with fewer than 3 oocysts.

**Figure S2:**
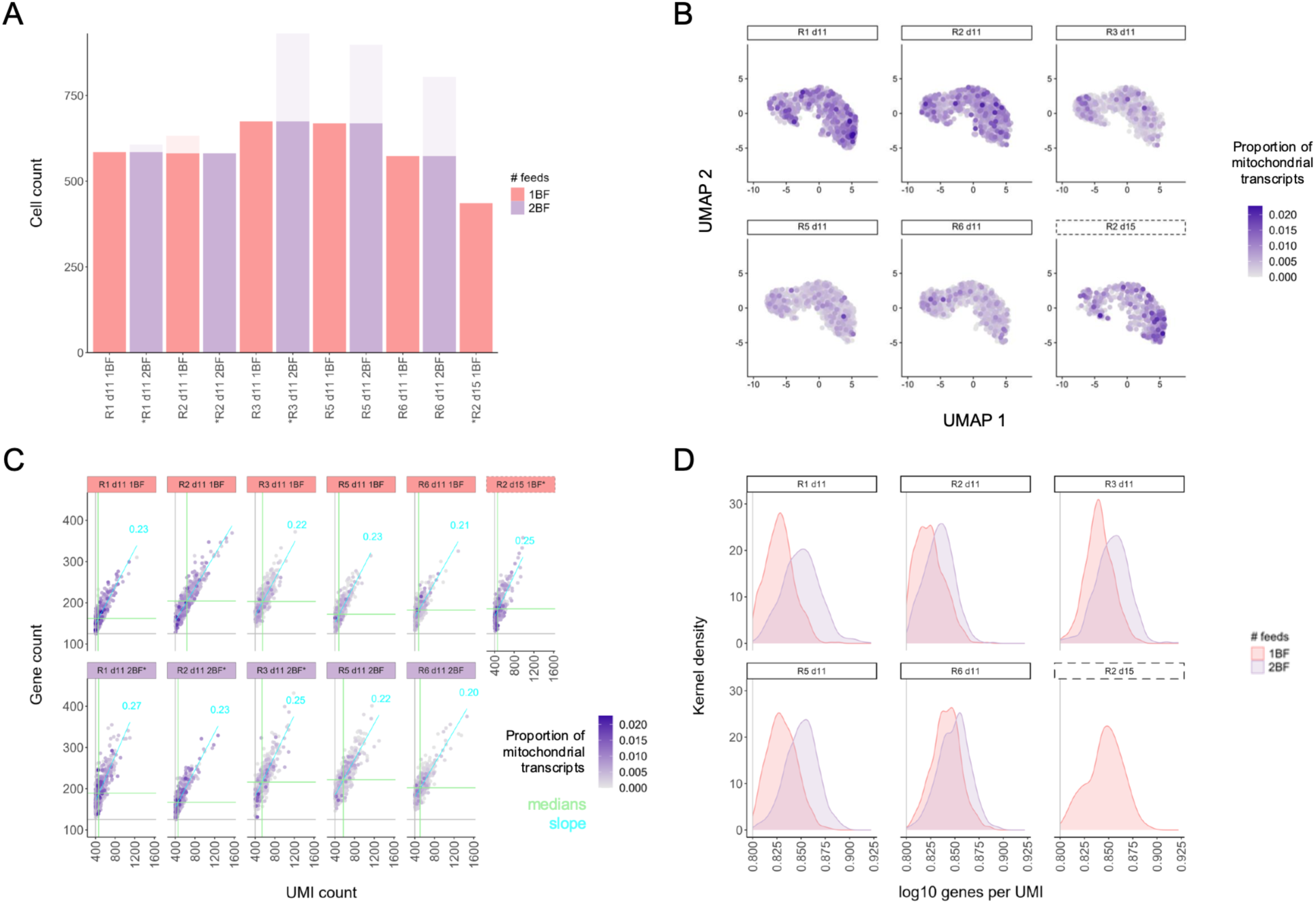
Quality control metrics for scRNA-seq data. (A) Cell counts per sample were randomly subsampled down to the minimum for each replicate (R1-R6). (B) All three clusters likely represent viable cells as mitochondrial gene representation does not exceed 0.025 of reads in any sample and does not correlate with UMAP clustering. The UMAP was generated *de novo* from raw UMI count matrices, incorporating both day 11 and day 15 data (rather than integrating day 15 data into an existing day 11 UMAP). (C) Gene counts and UMI counts are consistent across samples (axis medians in green) and correlate linearly (slope in blue). (D) Sequencing complexity (log_10_[genes per UMI]) is similar between samples and bloodfeeding regimens. The trend for higher complexity in 2BF samples may be biological, or technical due to dilution from higher sporozoite numbers. In (A) and (C), asterisks (*) indicate sporozoite samples derived from ds*GFP*-injected mosquitoes.

**Figure S3:**
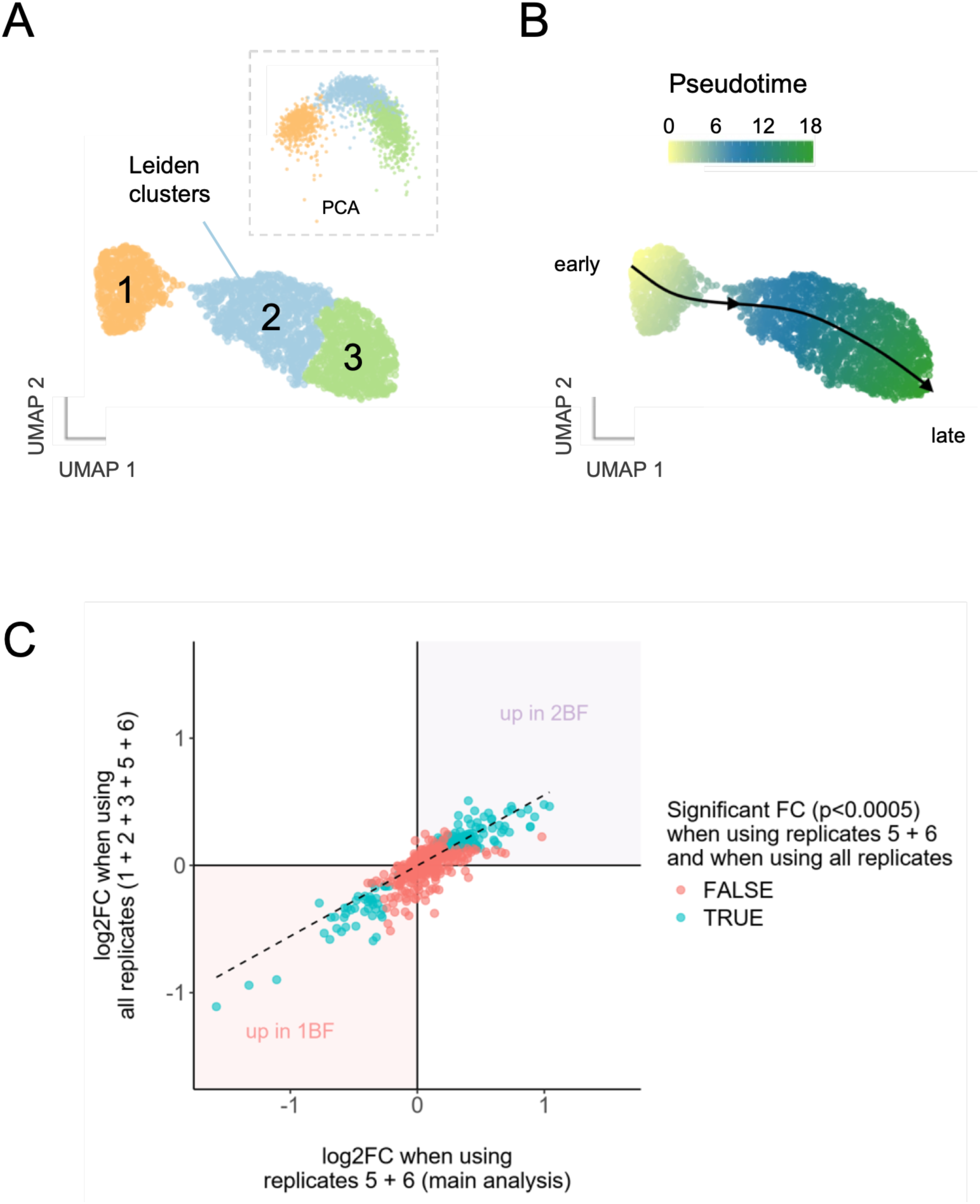
Analysis of scRNA-seq data focused on unmanipulated mosquitoes (replicates 5 and 6) is highly comparable to analyses pooling all scRNA-seq replicates (1–3,5 and 6). Re-running integration/reduction analysis gives similar clustering (A) and pseudotime (B) results to our primary analysis, which used all scRNA-seq experiments (1+2+3+5+6) to increase available anchoring points. (C) Results of our primary differential expression analyses, focused on unmanipulated mosquitoes with field-like infection intensities, are very similar to differential expression analyses that include all experimental replicates. Inclusion of the additional replicates in the ScranFindMarkers analysis dampens fold-change magnitude (slope <1), but 91.1% (down in 2BF) and 71.8 % (up in 2BF) of differentially expressed genes remain below the p-value threshold (p < 0.0005). See **Table S2**.

**Figure S4:**
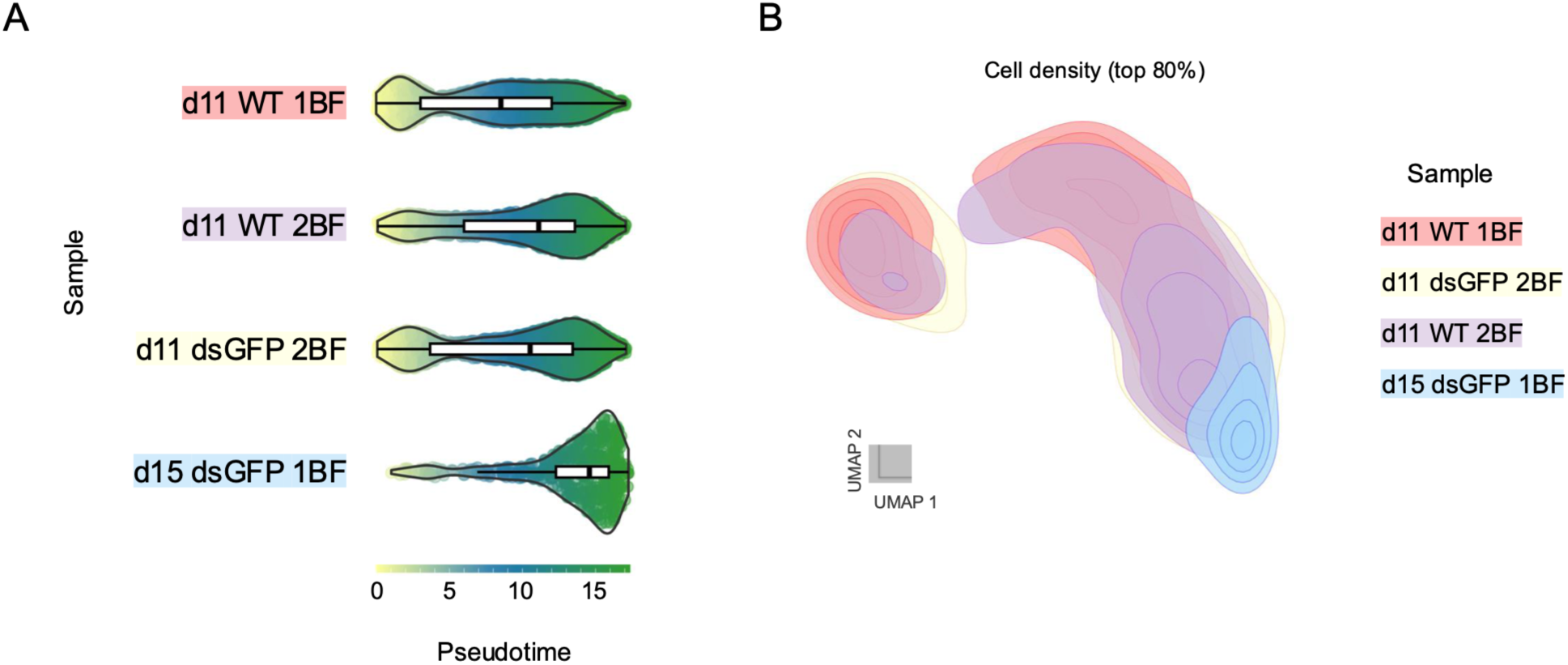
An additional blood meal shifts cellular abundances to a later pseudotime characteristic of more mature sporozoites. Directionality of shifts in cell abundance induced by an additional blood meal were confirmed towards mature sporozoites (right, dark green) through comparison to a single replicate of sporozoites harvested 15 d pIBM over pseudotime (A) and gene expression UMAP space (B). In (A), white box plots show median (central black line) and interquartile range of cellular pseudotime show a shift from WT 1BF sporozoites (5 replicates) to WT 2BF (2 replicates) and dsGFP 2BF sporozoites (3 replicates). In (B), kernel density plots incorporating day 11 and day 15 data highlight a shift away from top left to bottom right as sporozoites mature. Contours delimit areas containing the top 50% of cells and connect areas of similar cell density in 10% increments.

**Figure S5:**
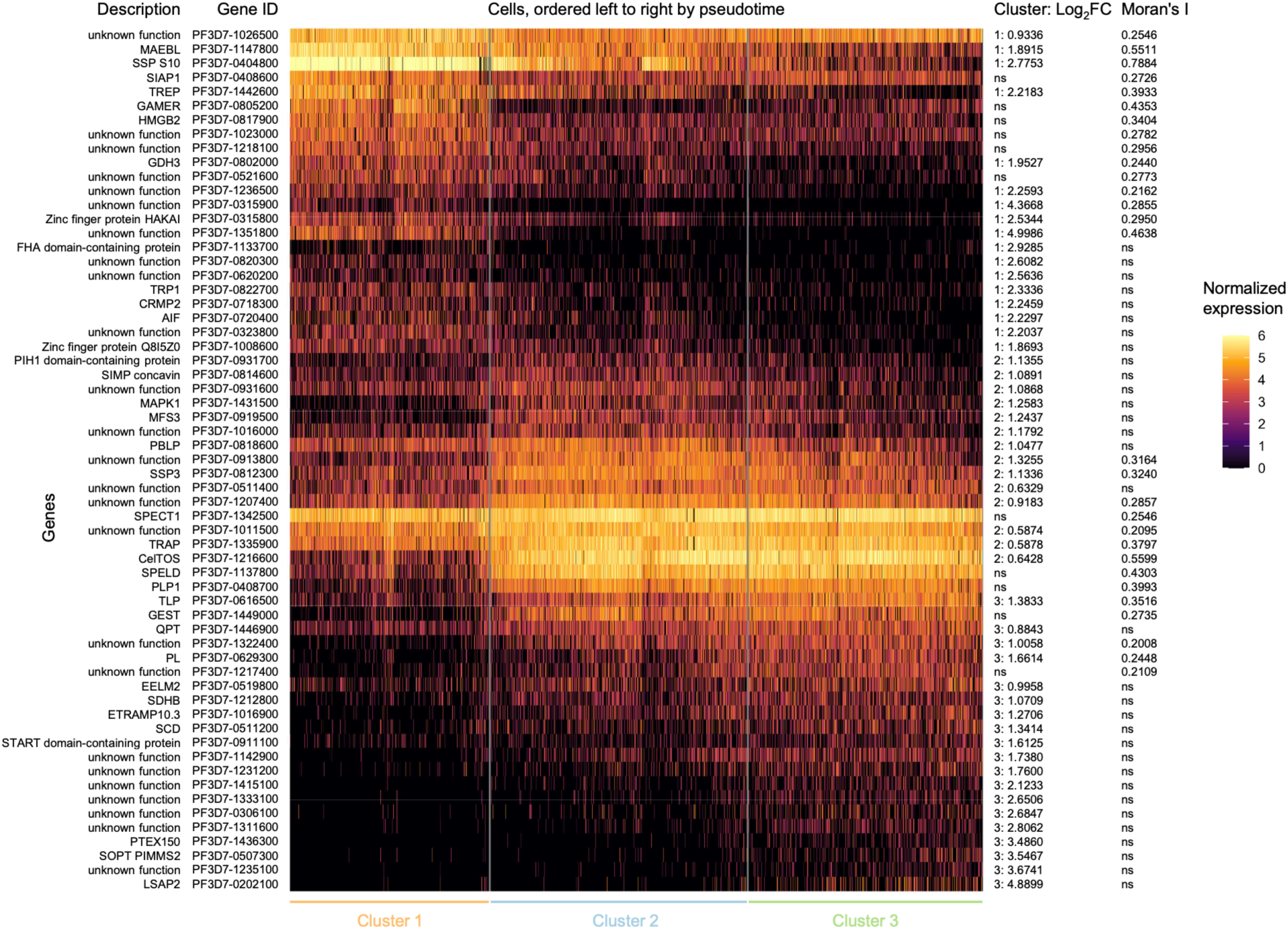
Expression heatmap for genes identified as primary cluster markers and/or showing autocorrelation in the UMAP space (Moran’s. **I)**. The heatmap includes all genes (rows) which were strongly supported as cluster markers (fold-change for member vs. non-member cells averaging 1.5 across the study’s 10 sample harvests, with the weakest p-value for any individual sample falling below 1e-4) based on Wilcoxon rank-sum testing with FindConservedMarkers in Seurat v5.0.3. Significant cluster associations and fold-change values are indicated in the penultimate column. The heatmap also includes all genes for which expression showed significant spatial autocorrelation signal based on Moran’s I > 0.20 (values in right column) and q-value < 1e-6 via ‘graph_test’ function in Monocle3 v1.3.1. Using discrete cluster comparisons vs. continuous coordinate analysis (respectively), these two complementary methods identify genes whose expression displays non-random patterning across the pseudotime trajectory. The analysis uses 1000 random 1BF sporozoites and 1000 random 2BF sporozoites after filtering for cells with >500 UMIs. These 2000 cells are ordered by pseudotime along the x-axis. Vertical divisions indicate Leiden cluster membership (labels at bottom). Cell fill color within each heatmap row represents normalized expression (color bar on the right).

**Figure S6:**
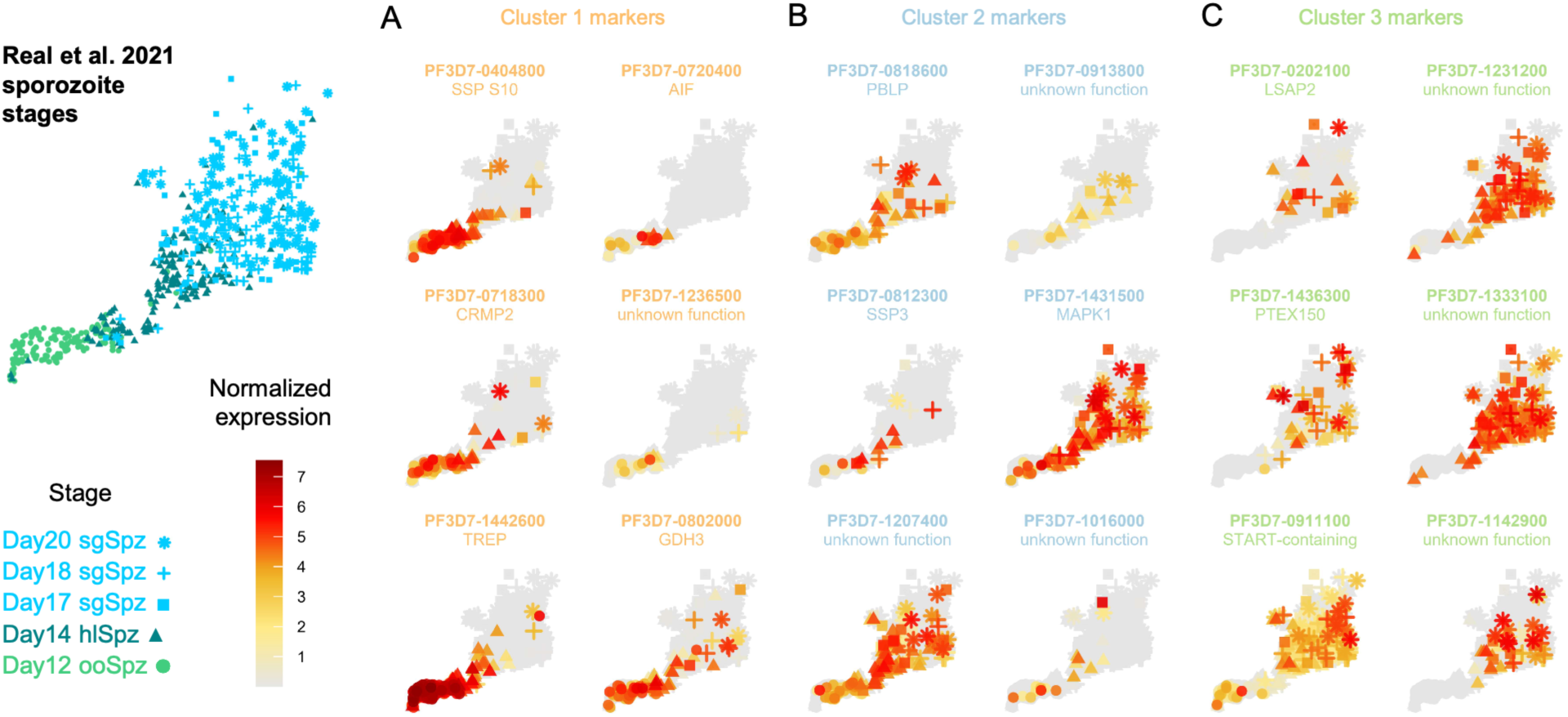
Expression of selected salivary gland sporozoite cluster marker genes in sporozoite stage data from the Malaria Cell Atlas (MCA) (**22**). Expression of cluster marker genes identified in this study of salivary gland sporozoites is shown on UMAPs plotting MCA sporozoite data (midgut, ooSpz, day 12); hemolymph, hlSpz, day 14; salivary glands, sgSpz, day 17–20). Genes marking salivary gland cluster 1 (A) overlap with previously identified ooSpz and hlSpz, whereas genes marking clusters 2 (B) and 3 (C) overlap with previously identified salivary gland sporozoites.

**Figure S7:**
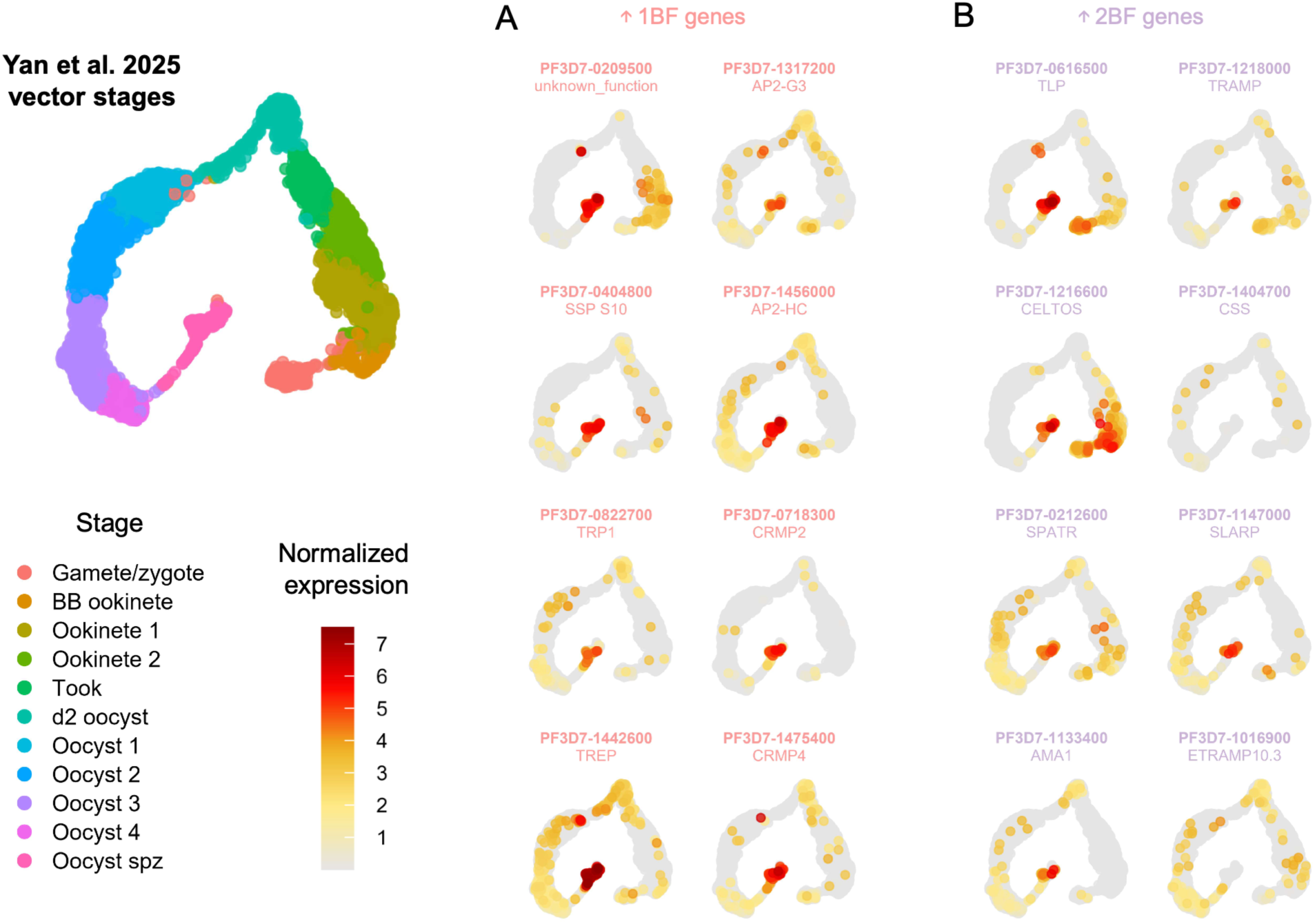
Differentially expressed genes show sporozoite-specific expression. UMAPs for selected genes downregulated (A) or upregulated (B) in 2BF sporozoites were generated using data from Yan, Verzier, Cheung et al. (40), Mohammed et al. (99), and the Malaria Cell Atlas (22). Selected genes are weakly expressed in earlier mosquito stages, and/or are expressed most strongly in oocyst sporozoites. The left panel shows how the UMAPs relate to the developmental stages of parasite development in mosquitoes.

**Figure S8:**
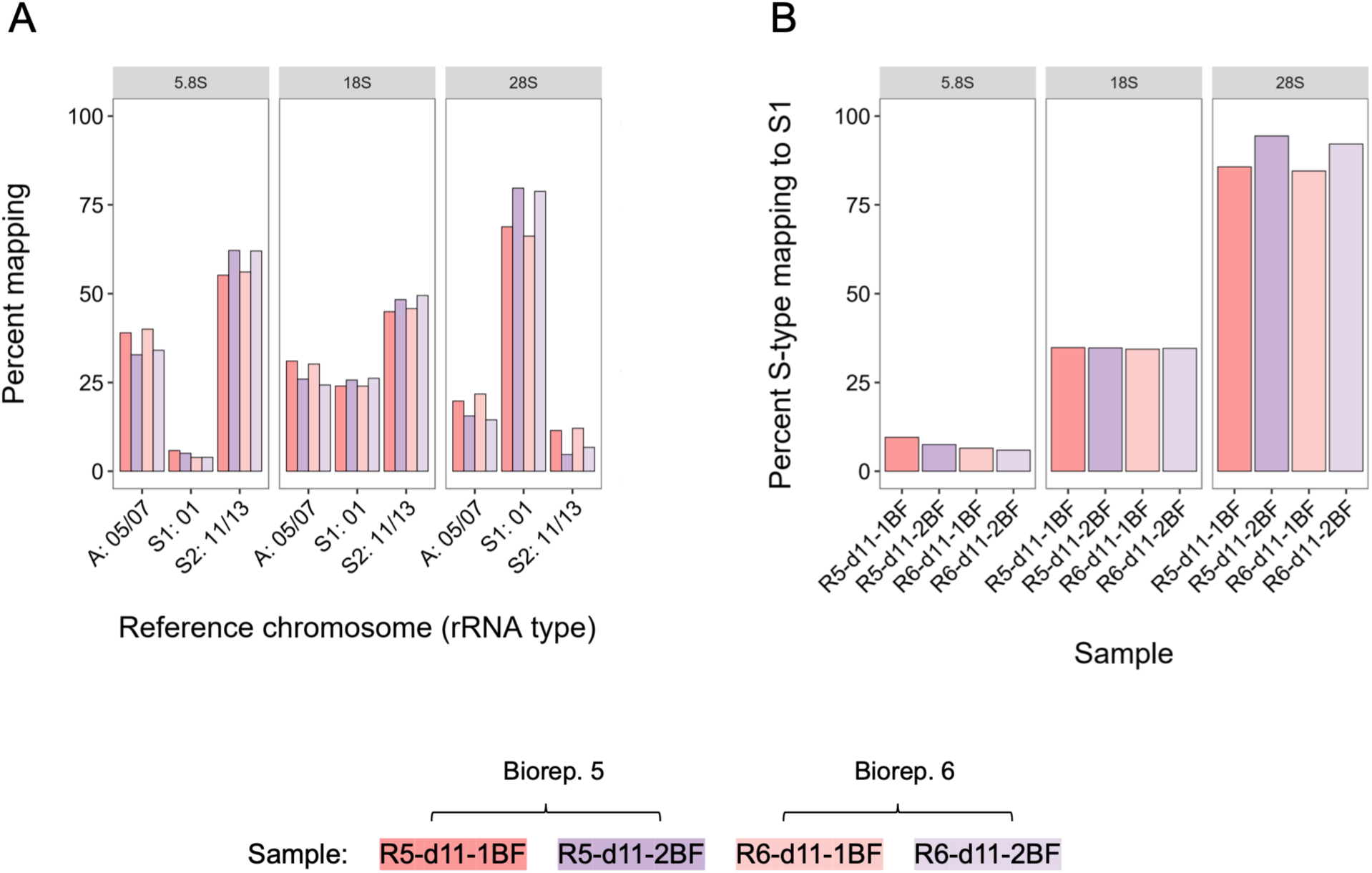
Percentage transcription and mapping of 5.8S, 18S and 28S ribosomal rRNA reads. Raw reads analyzed are from replicates 5 and 6 used in our primary differential gene expression analysis comparing 1BF and 2BF mosquitoes. A) Percentage of rRNA reads matching unambiguously to A-type (chromosomes 5 and 7), S1-type (chromosome 1) or S2-type (chromosomes 11 and 13) genomic loci in day 11 samples from 1BF and 2BF mosquitoes are shown, separated by ribosomal rRNA. After an additional blood meal (purple shades), sporozoites show a modest shift away from A-type rRNA loci towards S-type rRNA loci. B) Percentage of S-type rRNA reads that are S1-type reveals substantial divergence in abundance between rRNA genes, with most 5.8S and 18S expressed from S2-type loci, while most 28S are from S1-type loci. Note, inferences from raw reads without UMI counts requires caution.

**Figure S9:**
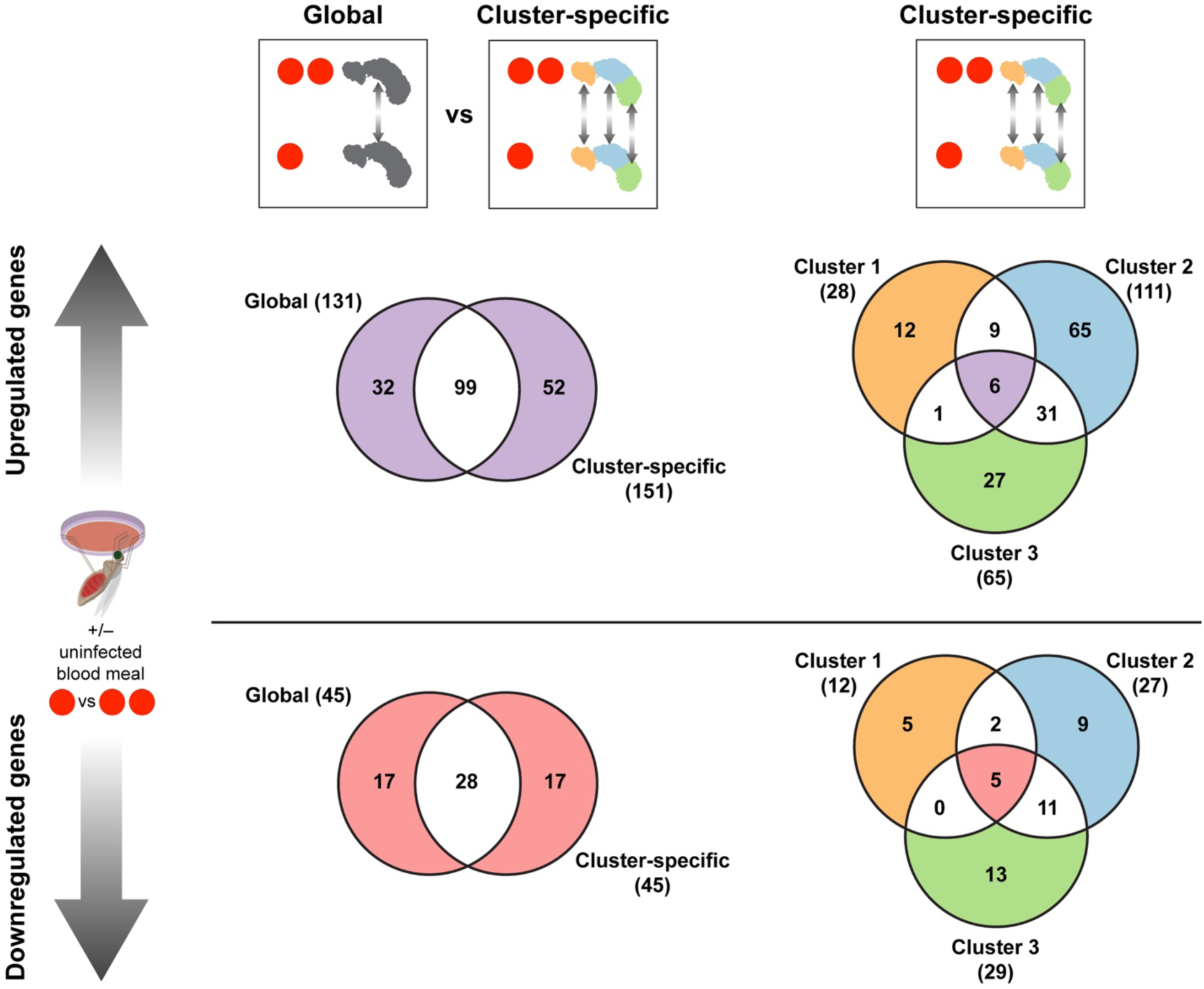
Overlap of differentially expressed gene lists between global and cluster-specific analyses. There is considerable overlap between upregulated (top) and downregulated (bottom) genes identified through global analysis (all cells) between 1BF (pink) and 2BF (purple) sporozoites to those identified when comparing cells within each Leiden cluster (c1, c2 and c3), suggesting differential abundance of mature sporozoites across clusters is not responsible for the observed transcriptional changes in our global analysis. Newly identified differentially expressed genes tended to be cluster specific, whereas those regulated in multiple clusters had previously been identified.

## Notes

### Competing Interest Statement

The authors have declared no competing interest.

